# Selective illumination on line-scanning confocal mesoscope enhances background rejection in cortex-wide mouse brain imaging

**DOI:** 10.1101/2024.05.29.596431

**Authors:** Hao Xie

## Abstract

Confocal microscope has optical sectioning that is accessible for structural and dynamic imaging in *in vivo* mouse brain. With the requirement of brain-wide field-of-view (FOV) in many neuroscience researches, existing confocal microscope fails to fulfill the requirement. Here, we proposed the brain-wide, high resolution line-scan confocal mesoscope (LSCM) for *in vivo* mouse imaging, achieving 6.6-mm-FOV, 3.2-μm-resolution and video-rate acquisition. To further enhance the background rejection ability, we introduced the selective illumination method into our system. We demonstrated that the proposed technique is s able to image the neurodynamics in *in vivo* mouse brain. Comparing to the LSCM and wide-field mesoscope, our selective illumination line-scan confocal mesoscope improves more than 25% and 3 times background rejection ability, respectively.

## 1. Introduction

The fluorescence microscope has been a preferred tool for structural and dynamic imaging in *in vivo* mouse brain [1]. With the neuroscience research progressing, recent studies reveal that variety cognitive processes involved the interaction of multiple brain regions, which are distributed across the entire mouse brain [2-4]. Thus, the brain-wide, cellular resolution mesoscope is required urgently in neuroscience field, especially with video-rate acquisition [5, 6]. Single-photon wide-field microscope, with advantages of wide field-of-view (FOV), diffraction-limit resolution and high frame rate, has been developed to a variety of brain-wide mesoscope. For example, the Firefly microscope [7] and COSMOS macroscope [8] have more than 6-mm FOV, diffraction-limit resolution at 30 Hz frame rate. MFIAS macroscope [9] and RUSH macroscope [10] demonstrate the ability of centimetre-wide FOV, micrometric resolution at video-rate acquisition, which are applicable to *in vivo* entire mouse brain imaging. While, wide-field illumination leads to severe background fluorescence, which degrades both contrast and resolution of the mouse brain image. To overcoming the issue, a variety of techniques have been proposed. Structured-illumination microscopy (SIM) [11, 12] synthesized several patterned-illuminating images to suppress the out-of-focus fluorescence, HiLo microscopy [13, 14] extracts in-focus information and enhances the image quality by combining a nonuniform illumination image and a uniform image for extracting in-focus information, deconvolution microscopy applied computational algorithms to remove out-of-focus fluorescence based on the point-spread-function of the system, photobleaching imprinting microscopy (PIM) [15] extracted high-order fluorescence signals from photobleaching-induced fluorescence decay to illuminate the out-of-focus fluorescence. While, these methods required several images to restructure high contrast image by computational algorithms, which are inevitable to induce artificial aberration and decrease frame-rate. Confocal microscopy [16, 17] by setting a small pinhole before the detector, and multi-photon scanning microscopy by natural nonlinear effect [18-20], have optical sectioning to reject the background fluorescence. While, due to point-scanning strategy, the frame rate was sacrificed inevitably compared to the wide-field microscope. Laser scanning confocal microscopy based on Mesolens system requires approximate 200 seconds for a full FOV (6 mm) [21]. 2-photon random access mesoscope (2p-RAM) has 1.9 Hz frame rate for 4.4 mm × 4.2 mm FOV [22], and Diesel2p mesoscope has 3.85 Hz for 5 mm × 3 mm FOV [23]. The limited imaging speed prevented their application in dynamic imaging. As a compromise strategy, line scanning confocal microscopy (LSCM) applied a narrow slit before the detector [24-26] or Rolling Shutter mode on camera [27]. Thus, LSCM could balance the frame-rate and out-of-focus rejection ability, which has been demonstrated in *in vivo* mouse brain. While, the existing LSCM did not extend FOV to brain-wide scale in mouse brain.

In this study, wo proposed a selective illumination line-scanning confocal mesoscope (SI-LSCM), achieving mouse brain imaging with brain-wide FOV, cellular resolution and video-rate acquisition. First, we utilized the Mesolens system to the line-scanning confocal technique, achieving 6.6-mm FOV, 3.2-μm resolution and video-rate acquisition. Considering the sparse distribution of neurons, selective illumination method was applied to only excite the neuron-regions based on prior image, further eliminating the background fluorescence. Finally, we demonstrated our technique was able to image the neurons in *in vivo* mouse brain.

## 2. Experimental set-up and imaging principle

A schematic of our SI-LSCM is shown in Fig. 1(a). The illumination beam is produced by a CW laser (MBL-III-473-100 mW, CNI), whose central wavelength is at 473 nm. The beam expander is applied to the laser beam to 9 mm. After reflected by a galvo mirror (GVS011/M, Thorlabs) and focused by a cylindrical lens, the laser beam is formed to line-shape on the a SLM (Spatial Light Modulator). A lens set, consisted of Lens1, Lens2 and Lens3, relays the beam to the Mesolens with 2x/0.5 NA (MVPLAPO 2 XC, Olympus). After passing through by the objective, the line-shape beam is focused to the mouse brain and scanned by the galvo mirror. The excited fluorescence is collected by the same Mesolens and passes through the dichroic mirror. Then, the tubelens with 1x/0.25 NA (MVPLAPO 1X, Olympus), used as tube lens, focuses the fluorescence to the sCMOS (pixel size: 6.5 μm, ORCA-flash4.0, Hamamatsu). A dual-band filter (wavelength: 520 ± 12.5 nm/630 ± 46.5 nm, #87-241, Edmund) is placed in the emission light path to eliminate the reflected laser light. The FOV of the system is approximate 6.6 mm, and each pixel in the sCMOS corresponds to 3.25 μm on the image plane.

**Fig. 1.**
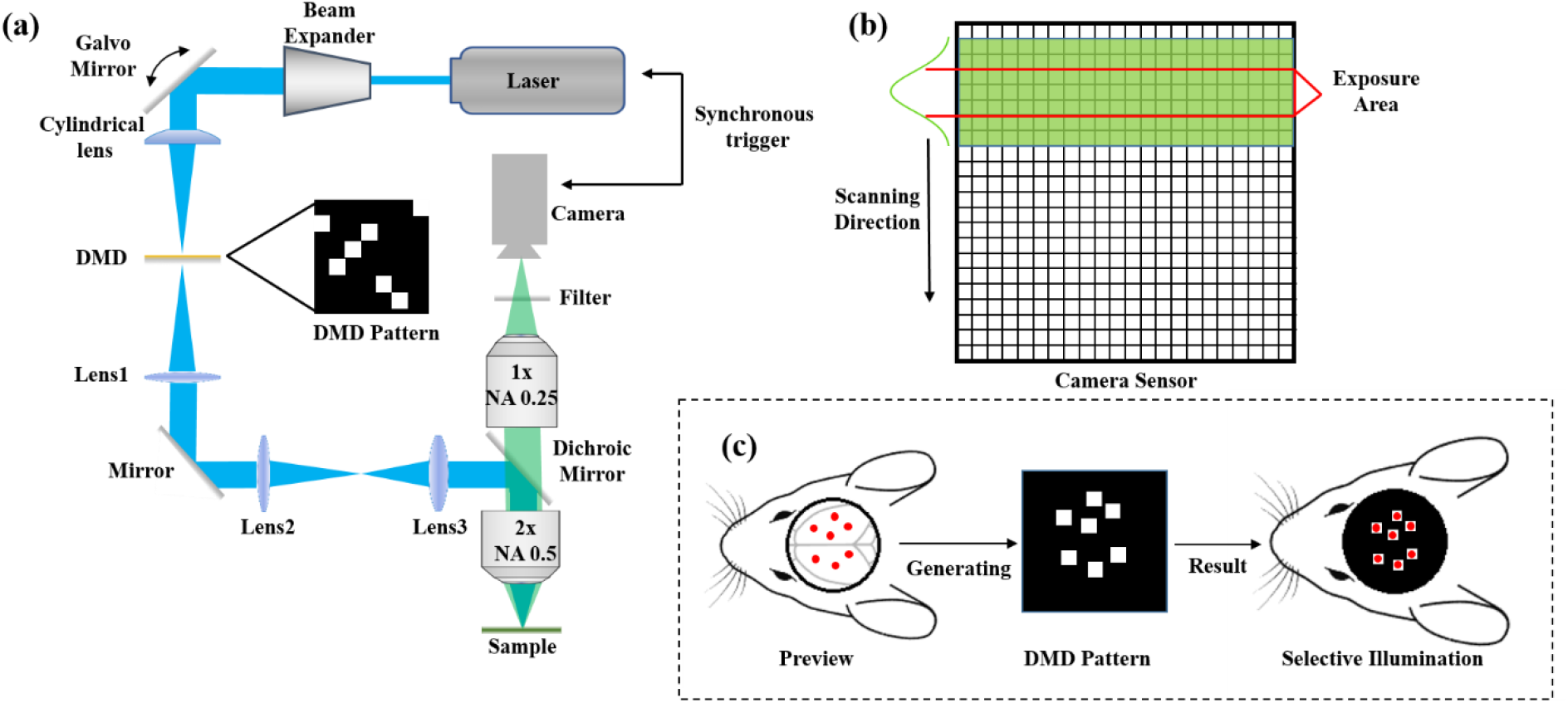
The principle of the selective illumination line-scanning confocal imaging. (a) The scheme of the custom-built SI-LSCM, with brain-wide FOV and cellular resolution. (b) Illustration of line-scanning signal recording method. (c) Schematic diagram of the selective illumination.

The confocal images were captured by the sCMOS with Rolling Shutter mode, which enables only several rows of the sensor to collect the signal. The Rolling Shutter mode can be used as adjustable slit. Fig. 1(b) illustrating the signal recording method, the sample-emitted line of fluorescence light has Gaussian distribution. Assuming the fluorescence light covers 7 rows and the Rolling Shutter slit can be set central 3 rows, the fluorescence on the other rows is be rejected by the slit. With the beam 1D scanning the whole FOV, the position of the slit moves synchronously and achieves the entire line-confocal image. Fig. 1(c) shows the principle of the selective illumination method, we firstly obtain 100 images of the mouse brain and ensure the positions of neurons. Based on the standard projection of neurons preview, the SLM pattern is then generated by the algorithm in Matlab (Math Works). Lastly, we project the designed pattern on the SLM, which only illuminate the neurons area.

## 3. Results

### 3.1 System Characterization

To evaluate the performance of our system, we firstly measured the beam width of excited line-shape focus on the scattered medium, whose intensity curve was shown in Fig. 2(a). Based on the FWHM (full widths at half maximum), the beam width is measured 226 μm. For optimizing the slit width of the system, we selected 25, 50, 100, 250 and 500 rows on the camera sensor to test the fluorescence decay. We measured the in-focus and out-of-focus fluorescence in the mouse brain. Fig .2(b) showed that the in-focus and out-of-focus fluorescence were decreased with the slit width decreasing. While, we set the slit width to 25 rows from 50 rows, the out-of-focus fluorescence was decreased less than the decrement of the in-focus fluorescence, resulting the signal-to-background ratio (SBR) degradation. Thus, we selected 50 rows as the slit width for our system. We further test the optical sectioning ability quantitatively by moving the 1-μm-diameter green fluorescent microsphere (T14792, Thermo Fisher Scientific) with different depths. The fluorescence energy of the wide-filed mode hardly changed with the depth change, but the fluorescence energy changed obviously for LSCM. The energy decreased at least 20% at ±50 μm depth, proving the sectioning ability of our LSCM. Lastly, one pixel of the DMD was corresponding to 70 μm on the focal plane, shown in Fig. 2(d).

**Fig. 2.**
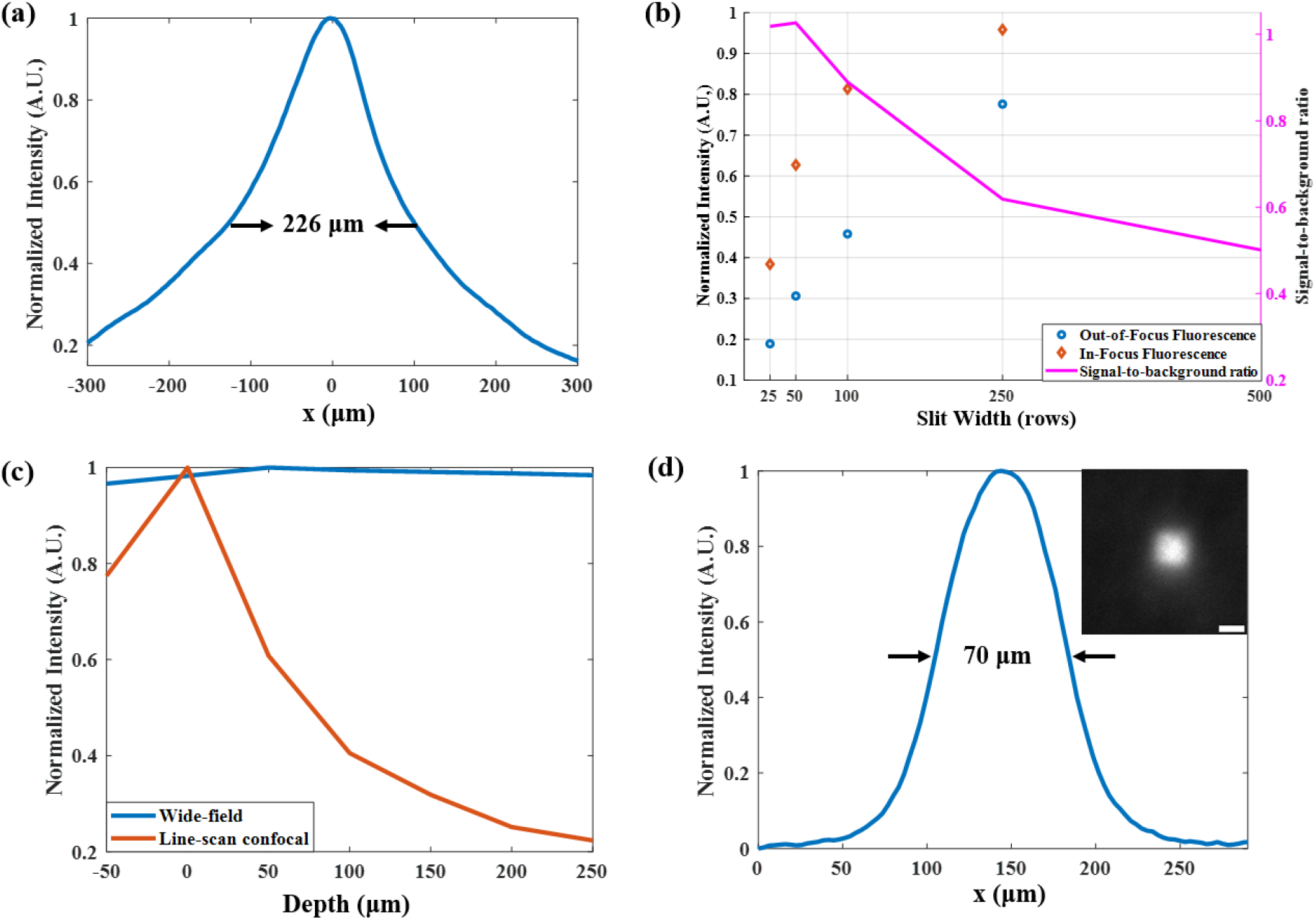
System Characterization of SL-LSCM. (a) Lateral intensity profiles of the line-shape beam. (b) In-focus and out-of-focus fluorescence with different slit widths. (c) Fluorescence energy with different depths. (d) Lateral intensity profiles of one pixel of the DMD on focal plane. Scale bar: 60 μm.

**Fig. 3.**
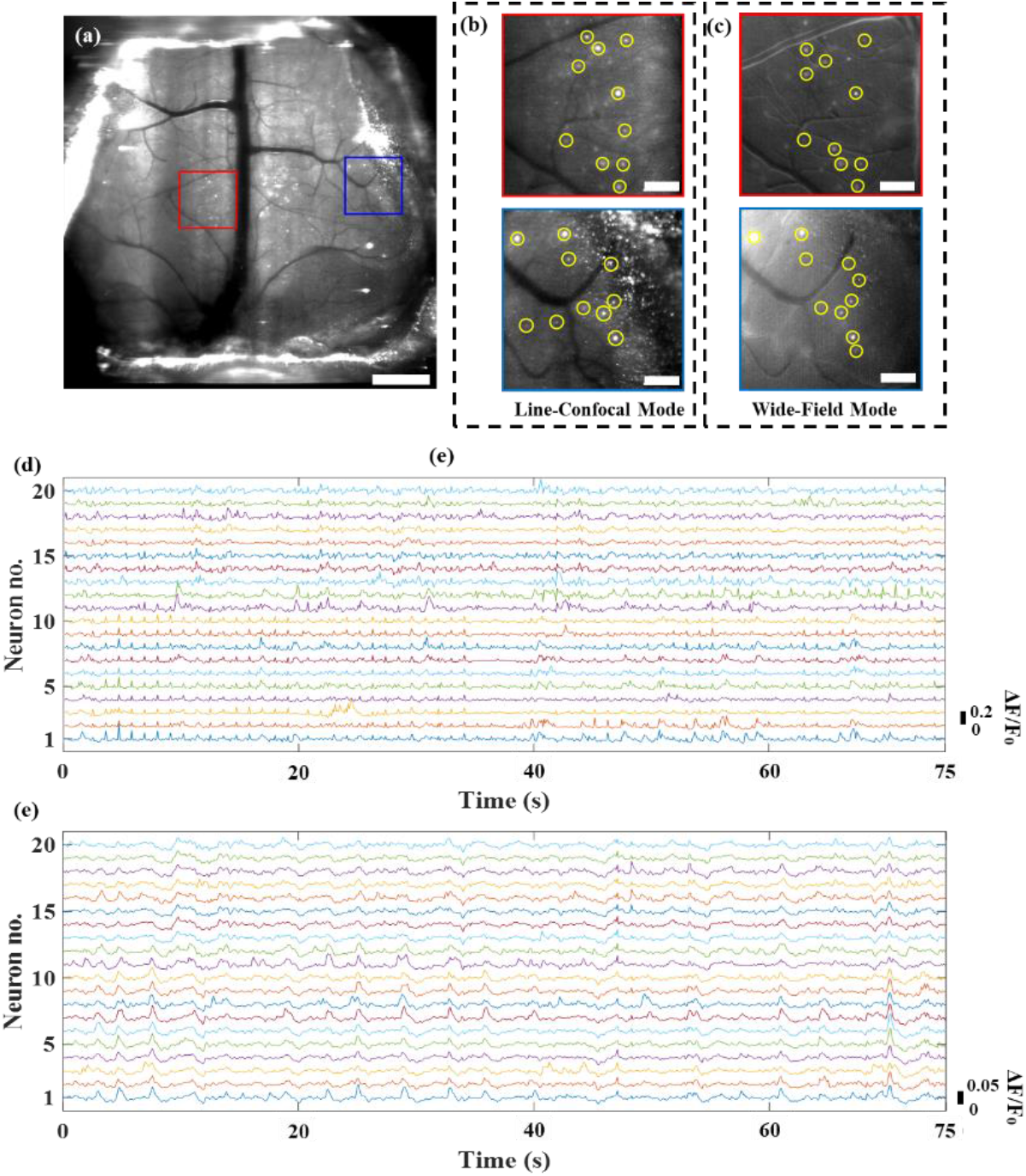
In vivo dynamic neuronal imaging in mouse brain. (a) Maximum deviation projection of 750 time-lapse images by Line-Confocal mode. Scale bar: 1 mm. (b) Magnified views of the subregions marked in (a). (c) The same subregions captured by Wide-Field Mode. Scale bar: 200 μm. (d, e) Fluorescence signal fluctuation (ΔF/F_0_) of neurons in yellow circles marked in (b, c).

### 3.2 In vivo mouse brain imaging by brain-wide LSCM

We then showed that our system enables *in vivo* dynamic neuro-imaging in Ai148 (TIT2L-GC6f-ICL-tTA2)-D;Rasgrf2-2A-dCre (JAX 030328, JAX 022864) mouse brains. Line-Confocal mode was achieved by setting 50 row of the camera as slit and frame rate as 10 Hz. Fig3. (a) showed the maximum deviation projections of 750 time-lapse images, performing 6.6 mm FOV and cellular-level resolution. To better show the ability of our system, a central and side subregions were selected and magnified in Fig.3 (b), the neurons in both subregions were identified clearly. We selected 10 neurons in each subregion and plotted the fluorescence signal fluctuation (ΔF/F_0_) curves, shown in Fig.3 (d). The maximum ΔF/F_0_ of the spikes was 0.28 and average ΔF/F_0_ was 0.157. To comparing our brain-wide line-scanning confocal mesoscope and wide-field mesoscope, we switched our system to Wide-field Mode by setting the exposure mode to global shutter and frame rate as 10 Hz. Fig.3 (c) showed the subregions and selected neurons captured by Wide-field Mode, and Fig.3 (e) plotted the ΔF/F curves of the neurons. The maximum ΔF/F_0_ of the spikes was 0.08 and average ΔF/F_0_ was 0.052. The results indicated that our system not only achieved brain-wide FOV and cellular-level resolution, but also enhance 3-times background rejection ability compared to wide-field mesoscope。

### 3.3 In vivo mouse brain imaging by brain-wide SI-LSCM

To further improve the background rejection ability, we introduced the selective illumination method into the system our brain-wide LSCM. The pattern on the SLM was calculated based on 100 time-lapse images. Fig. 4(a) showed the maximum deviation projections of 1000 time-lapse Ai148-D;Rasgrf2-2A-dCre mouse brain images captured by SI-LSCM, 55 neurons were labeled by red circles for plotting the fluorescence signal fluctuation (ΔF/F_0_) curves, shown in Fig. 4(d). The maximum ΔF/F_0_ of the spikes was 0.679 and average ΔF/F_0_ was 0.31. To better present the background rejection enhancement of SI-LSCM, we compared the SI-LSCM to the LSCM and the selective illumination wide-field mesoscope (SI-WF) with the same neurons and frame rate. Fig. 4 (e) and Fig. 4 (f) plotted the fluorescence signal fluctuation (ΔF/F_0_) curves captured by the LSCM and SI-WF, respectively. For the LSCM, the maximum ΔF/F_0_ was 0.384 and average ΔF/F_0_ was 0.212, and for the SI-WF, the maximum and average value were 0.283 and 0.166. Statistical results were presented in Fig. 4(g). Although combining with selective illumination method, the background rejection ability of the SI-WF could not reach the effect of the LSCM. While, the results demonstrated that the SI-LSCM had better background rejection ability than the LSCM, and the effect of background rejection improved 50%.

**Fig. 4.**
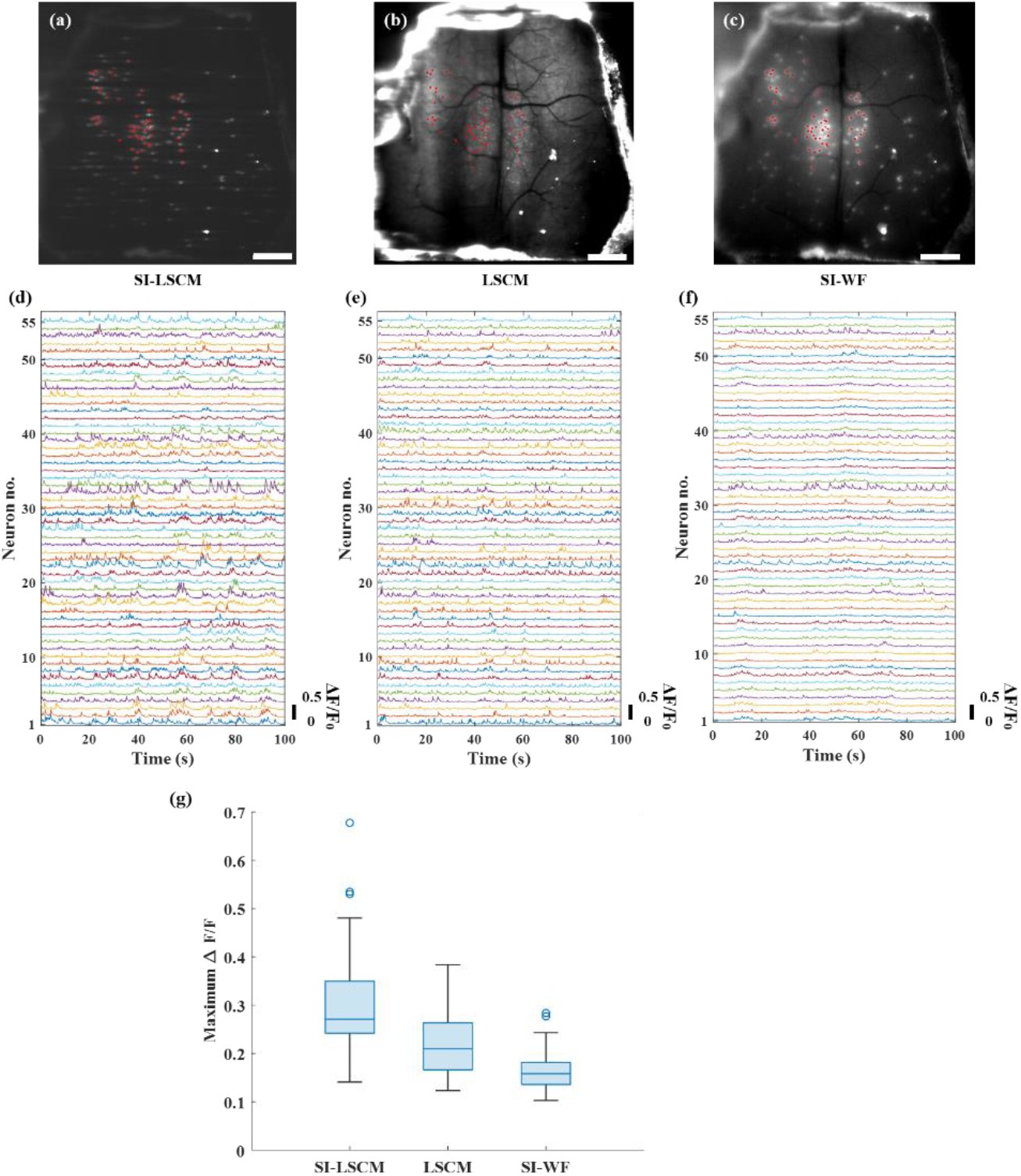
Comparison of SI-LSCM, LSCM and SI-WF in *in vivo* dynamic neuronal imaging. (a, b, c) Maximum deviation projection of 150 time-lapse images by SI-LSCM, LSCM and SI-WF. Scale bar: 1 mm. (d, e, f) Fluorescence signal fluctuation (ΔF/F_0_) of neurons marked by red circles in (a, b, c). (g) Statistical results of the maximum ΔF/F_0_ captured by SI-LSCM, LSCM and SI-WF.

## 4. Conclusion

In summary, we have demonstrated our brain-wide SI-LSCM for structural and functional imaging in mouse brain. The abilities of 6.6 mm FOV and 3.25 μm resolution allow the SI-LSCM to observe neurodynamics distributed across entire mouse brain. The selective illumination method is utilized reject background fluorescence and improved contrast by fully using the properties of line-scanning confocal microscopy. Through the experiments, the results demonstrated that the background rejected of the SI-LSCM enhances more than 3 times than the wide-field mesoscope and 50% than the LSCM with the same conditions, proving the effect and practicability of our technique.

## Author Contributions

Hao Xie supervised this research, conceived and designed the project. Chaowei Zhuang designed and built the system, analyzed the results, and wrote the manuscript.

